# Antibacterial activity of human natural killer cells in the absence of accessory cells against extracellular *Staphylococcus aureus* and hypervirulent *Pseudomonas aeruginosa* bacteria

**DOI:** 10.1101/2023.11.13.566836

**Authors:** Audrey Guilbaud, Baptiste Lottin, Gwenann Cadiou, Tiffany Beauvais, Sylvia Lambot, Barbara Mouratou, Nathalie Labarrière, François Davodeau, Frédéric Pecorari

## Abstract

Natural killer (NK) cells play a crucial role in the innate immune response to bacterial infections, including those due to *Pseudomonas aeruginosa* (*P. aeruginosa*) and *Staphylococcus aureus* (*S. aureus*). *In vivo*, it has been shown that NK cells are activated by innate accessory cells that detect the presence of bacteria and activate NK cells via a cytokine network. *In vitro*, several studies have shown that NK cells can also be activated without the help of accessory cells by direct contact with some bacteria species such as extracellular *P. aeruginosa*. Whether this phenomenon of direct activation is restricted to certain bacterial species, or whether it can be generalized, is still debated, as for example in the case of NK cell activation by *S. aureus*, which seems to require the intervention of accessory immune cells. Here, we show with co-incubation experiments between NK cells and two bacterial species, that, in the absence of accessory cells, NK cells are able to impede bacterial growth. This has been demonstrated for the *P. aeruginosa* PA14 strain, which is hypervirulent and known for its deleterious effects on NK cells, as well as for the *S. aureus* Newman strain. The monitoring of CD107a by flow cytometry suggests that NK cells degranulate after contact with *S. aureus* bacteria. Our study contributes to the idea that NK cells can be activated in the absence of any accessory cells by various species of bacteria, even an hypervirulent one, and therefore that NKs can directly have an antibacterial effect. This important insight may pave the way for new therapeutic approaches using antibacterial NK-cell engagers.

## INTRODUCTION

NK cells are a type of lymphocyte that plays a crucial role in the innate immune system. They are mostly known for their ability to target and eliminate virus-infected cells and cancer cells. However, the role of NK cells in bacterial infection has been described historically since the 1980s, with their ability to lyse cells infected with *Shigella flexneri*, *Legionella pneumophila* or *Mycobacterium avium* [1–3], and more recently for intracellular *Listeria monocytogenes* [4]. Although the crucial role of NK cells in fighting bacterial infections is quite well documented, some aspects of their activation by bacteria are still partly described. A mechanism of NK activation which has been the most studied, is based on recognition of pathogen-associated molecular patterns (PAMPs) by pathogen-recognition receptors (PRRs) at the surface of accessory cells, such as macrophages, neutrophiles, dendritic cells, and B lymphocytes that detect the presence of bacteria. These activated accessory cells then secrete cytokines, notably IL-15 and IL-18, that play an important role in the development, proliferation, survival and activation of NK cells [5]. NK cell activation is characterized by the expression of CD69 and interferon gamma (IFN-γ) and leads to the polarized release of granules with a cytotoxic lytic effect, thanks to perforin, granulysin and the granzymes they contain. These effectors are able to induce significant physical damage to bacterial walls, leading to their death [6].

About two decades ago, the possibility of NK cells being activated by another mechanism *via* direct contact with bacteria has been questioned [7]. Although increasing evidence show that NK cells can have antibacterial activity independently of accessory cells, this has long remained open to doubt as it contradicts the paradigm that NK cells play their role under the control of a network of signals relayed by accessory cells [8]. However, several studies have reported that NKs can be activated by contact, for instance with mycobateria, *Nocardia farcinica*, *Pseudomonas aeruginosa*, *Klebsiella pneumoniae*, *Escherichia coli*, *Salmonella typhimurium*, *Staphylococcus epidermidis* and *Burkholderia cepacia* [7, 9–13], thereby inducing an antibacterial effect. Thus, even in the absence of accessory cells, a broad spectrum of bacteria can activate NKs, including coccus, bacilli, Gram-negative and Gram-positive bacteria. When studied at the molecular level, bacterial recognition can involve a natural cytotoxicity receptor (NCR) located on the NK surface [11].

This broad potential of NKs to kill a variety of bacteria could provide the basis for the development of innovative therapeutic approaches, irrespective of the antibiotic resistance status of the targeted pathogens. Despite the introduction of more than 30 new antibiotics since 2000, the supply chains that generate new drugs are shrinking for all pathogens [14]. The emergence of antibiotic-resistant strains is far from being under control, according to the world health organization (WHO), and cases are multiplying in many countries of diseases for which there are no longer any non-invalidating treatments. In 2008, *Enterococcus faecium*, *S. aureus*, *Klebsiella pneumoniae*, *Acinetobacter baumannii*, *P. aeruginosa* and *Enterobacter* species were grouped under the ESKAPE acronym as pathogens known to escape the lethal action of antibiotics [15]. In 2017, the WHO has published the priority pathogen list (PPL) for R&D of new antibiotics that includes most of those from the ESKAPE group, plus other species such as *Helicobacter pylori* and *Campylobacter*[16]. For most of these pathogens, multidrug resistant clinical isolates are found, and they are the leading cause of nosocomial infections.

Multidrug-resistant (MDR), extensively drug-resistant (XDR) and Pan-resistant (PDR) strains, including *P. aeruginosa*, have been defined according to their non-susceptibility profile to various categories of antibiotics, the PDR strains being defined as non-susceptible to all agents in all antimicrobial categories [17, 18]. *P. aeruginosa* is a Gram-negative bacillus and an opportunistic pathogen often involved in nosocomial infections, most notably in patients with cystic fibrosis or those in intensive care units [17]. Upon infection of mice by *P. aeruginosa*, NKG2D expression is increased and NK cells participate to the clearance of bacteria [19, 20]. Furthermore, the depletion of NK cells leads to increased mortality rate of infected mice [21]*. In vitro,* human NK cells have been shown to bind directly to *P. aeruginosa* via their NKp44 receptor [11]. It was also demonstrated that human NK cells could exert a microbicide effect via granzymes on extracellular bacteria in the absence of any accessory cells [13].

*S. aureus* is also an opportunistic pathogen responsible for a large number of hospital-acquired infections and community-acquired infections. This Gram-positive coccus can cause numerous diseases such as skin infections, sepsis, pneumonia, infective endocarditis and bloodstream infections [22]. During the last decades, the spreading of MDR and XDR strains have been observed worldwide. Methicillin-resistant *S. aureus* (MRSA) strains are of concern as they are responsible for infections more difficult to treat, requiring the use of non-beta-lactam antibiotics, such as vancomycin or daptomycin. However, the increased use of vancomycin as a treatment of last resort has led to the emergence of vancomycin-resistant strains of *S. aureus* (VRSA), which poses a very serious threat to human health. The crucial role of NK cells in the immune response during infection by *S. aureus* was demonstrated with NK-depleted mouse models and IL-15 knockout mice [23]. The absence of NK cells makes mice susceptible to *S. aureus* infection, while IL-15 has been shown to have an indispensable effect on NK cells that cannot be compensated for by other immune cells. *In vitro*, it was reported that NK cells in presence of *S. aureus* bacteria require a co-stimulation by monocytes to be activated as assessed by CD69 and IFN-γ levels. A direct contact between *S. aureus* and NK cells was inefficient to activate NKs and no bacterial viability tests were performed [24]. Surprisingly, a previous study reported for another *Staphylococcus*, *Staphylococcus epidermidis* (*S. epidermidis*), that NK cells were sufficient to show a strong antibacterial effect *in vitro* in a contact-dependent manner according to counting of survival bacteria [12]. Thus, NK cells play a critical role in showing an antibacterial effect, but it seems that depending on the bacterial species, co-stimulation by accessory cells is either mandatory or not.

The main objective of this work was to determine to which extent NK cells, in the absence of accessory cells, are able to interfere with pathogenic bacteria growth, whether bacteria are Gram-bacilli or Gram+ coccus. To this end, in absence of accessory cells, we co-incubated NK cells with the hypervirulent *P. aeruginosa* PA14 strain and the *S. aureus* Newman strain, both belonging to the ESKAPE group. The antibacterial activity of NKs was studied at different NK/bacteria ratios and as a function of incubation time, by determining bacterial survival by counting colony forming units on petri dishes and by studying NK degranulation using flow cytometry.

## MATERIALS AND METHODS

### Bacteria strains

The *S. aureus* Newman strain and the *P. aeruginosa* PA14 strain were kindly provided by Prof. Timothy Foster (Dublin, Ireland) and by Dr. Alexis Broquet (Nantes, France), respectively.

### NK cell line

The human NK92-CD16 cell line was provided by Dr. Henrie Vié [25] and was cultured in RPMI 1640 medium with 10% FBS (Eurobio), 2 mM GlutaMAX-I (Gibco), 100 U/mL penicillin (Gibco) and 0.1 mg/mL streptomycin (Gibco).

### Sorting and amplification of NK

PBMC were isolated from heparinized blood from healthy adult volunteers by gradient centrifugation on Ficoll (Lymphocytes separation medium, Eurobio). All blood donors were recruited at the Blood Transfusion Center (EFS, France) and informed consent was obtained from all individuals. NK cells were collected by magnetic cell sorting using the EasySep Human NK Cell Isolation Kit (Stem Cell, France) according to the manufacturer’s instructions.

The purity of each NK population was checked before and after cell sorting and routinely by flow cytometry with anti-human CD16-BV421 (clone 3G8, BD Pharmagen, France), anti-human CD56-PeCy7 (clone B159, BD Pharmagen, France), anti-human CD3-FITC (clone BW264/56, Miltenyi Biotech, France).

After cell sorting, NK cells were expanded on allogeneic irradiated feeder cells (peripheral blood mononuclear cells and B-EBV B cells) with 150 U/mL human rhIL-2. Briefly, 2000 NK cells/well were distributed in 96-well round bottom plates with 200 μl of RPMI culture medium containing 8% human serum (Eurobio, France), 2 mM GlutaMAX (Gibco, France), 100 U/mL penicillin (Gibco, France), 0.1 mg/mL streptomycin (Gibco, France), human rIL-2 150 U/ml (Proleukin, Novartis) and irradiated EBV-B feeder cells (1.10^3^/well), and allogeneic PBMC from two donors (1.10^4^/well). The culture medium was changed every 2 days. NK cells were split when their number exceeded 1,5.10^5^/well.

### Killing Assay

An isolated colony of *P. aeruginosa* or *S. aureus* was picked from a Luria-Bertani (LB)/agar petri dish to inoculate 10 mL of LB medium that were incubated 16 h at 37°C with shaking. Then 1 mL of the culture was used to inoculate 10 mL of LB medium and the culture was incubated for 2 h at 37°C with shaking. The culture was diluted in RPMI medium to distribute 5.10^2^/well bacteria per well in 96 well round bottom plates. Two days before the killing assay, NK cells were cultured in RPMI 1640 medium with 10% FBS (Eurobio), 2 mM GlutaMAX-I (Gibco) without antibiotics. The day of the killing assay, NK cells were washed 3 times in serum and antibiotic free RPMI 1640 media to discard any traces of antibiotics. Bacteria were co-cultured with or without NK cells (NK92hCD16 or amplified NK – from 2,5.10^2^/well to 5.10^4^/well) in quadruplicate in serum and antibiotic free RPMI 1640 media. Cultures were incubated at 37°C and 5% CO2 for 2 to 6 h. Co-cultures were diluted in sterile water and 10 µL were spotted on LB agar plate. Plates were incubated overnight at 37°C and the colonies were counted and recorded.

### Degranulation Assay

NK92hCD16 or amplified NK (2.10^5^/well) were stimulated for 4 hours with or without *S. aureus* Newman (from 2.10^6^ to 2.10^4^/well) in the presence of anti-human CD107a-AF647 mAb (clone H4B4, Biolegend, France) added at the beginning of stimulation in serum and antibiotic free RPMI media in 96 well round bottom plates. Cultures were incubated at 37°C and 5% CO2 for 4 h. To detect NK, cells were incubated with anti-human CD56 PE (clone B159, BD Pharmagen, France). Cells were next fixed with PBS 4% paraformaldehyde for 15 minutes. Stained cells were acquired on a BD FACSCanto II flow cytometer using DIVA software.

### Statistics

Statistical analyses were performed using GraphPad Prism 8 software. One-way ANOVA with Bonferroni correction was used to evaluate statistical significance, a p-value of less than 0.05 was considered statistically significant (*), p < 0.01 (**), p < 0.001 (***) and p < 0.0001 (****) as highly significant. For non-significant results, (ns) was indicated.

## RESULTS

### NK cells from healthy donors impede the growth of an hypervirulent strain of *P. aeruginosa*

We first studied the efficacy of NKs against a *P. aeruginosa* strain different from the PAO1 strain studied in the work of Feehan et al [13]. We chose the PA14 strain with the aim to challenge antibacterial efficacity of NK cells, this strain being described as hypervirulent compared to PAO1 [26], and also because PA14 was reported to mediate NK cell apoptosis *in vitro* [27].

NK cells were isolated from healthy donors as described in “Materials and methods” section. Flow cytometry measurements (figure 1) showed that these NK cell populations were highly pure, around 93%.

**Figure 1:**
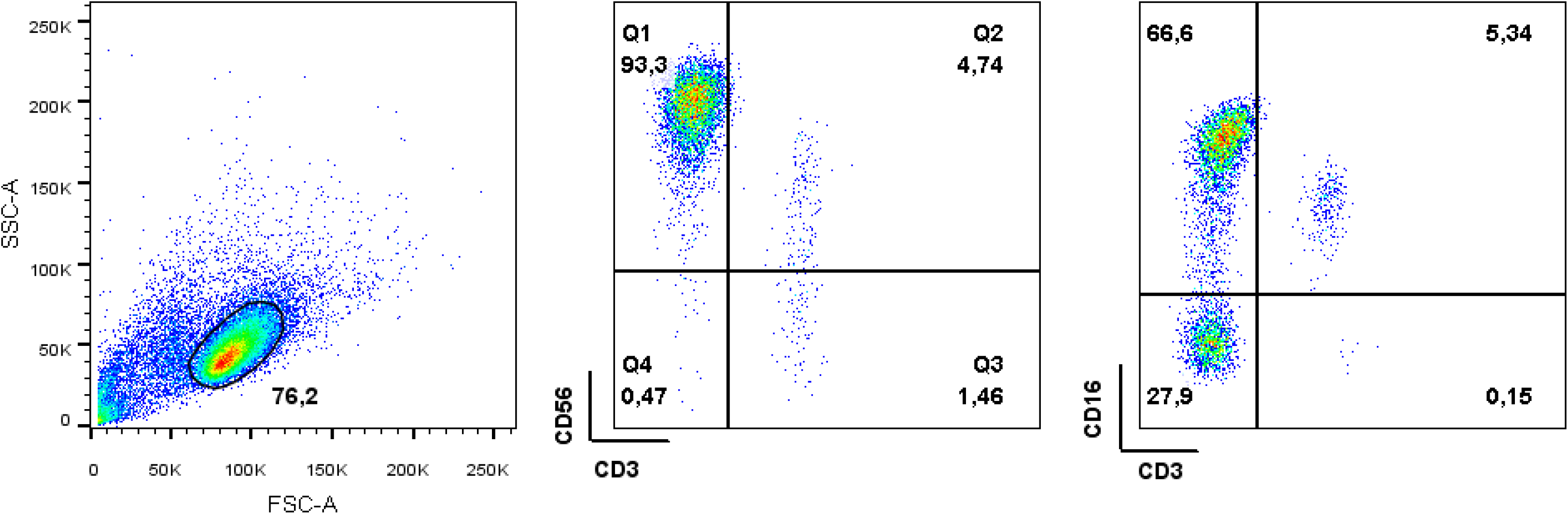
Control of NK cell purity from healthy donors. NK cells derived from healthy donor D3 were stained with FITC-conjugated anti-CD3, PECy7-conjugated anti-CD56 and BV421-conjugated anti-CD16 to control purity.

We then set up co-incubations with a constant number of bacteria per condition (500 CFU) and an increasing number of human NK cells to study the effect of effector:target (E:T) ratio on *P. aeruginosa*. The figure 2A shows that after 6h of co-incubation, PA14 CFU numbers were reduced as the E:T ratio increased compared to the condition without NK cells. The effect on bacterial growth was already optimal at 4 h (figure 2B).

**Figure 2:**
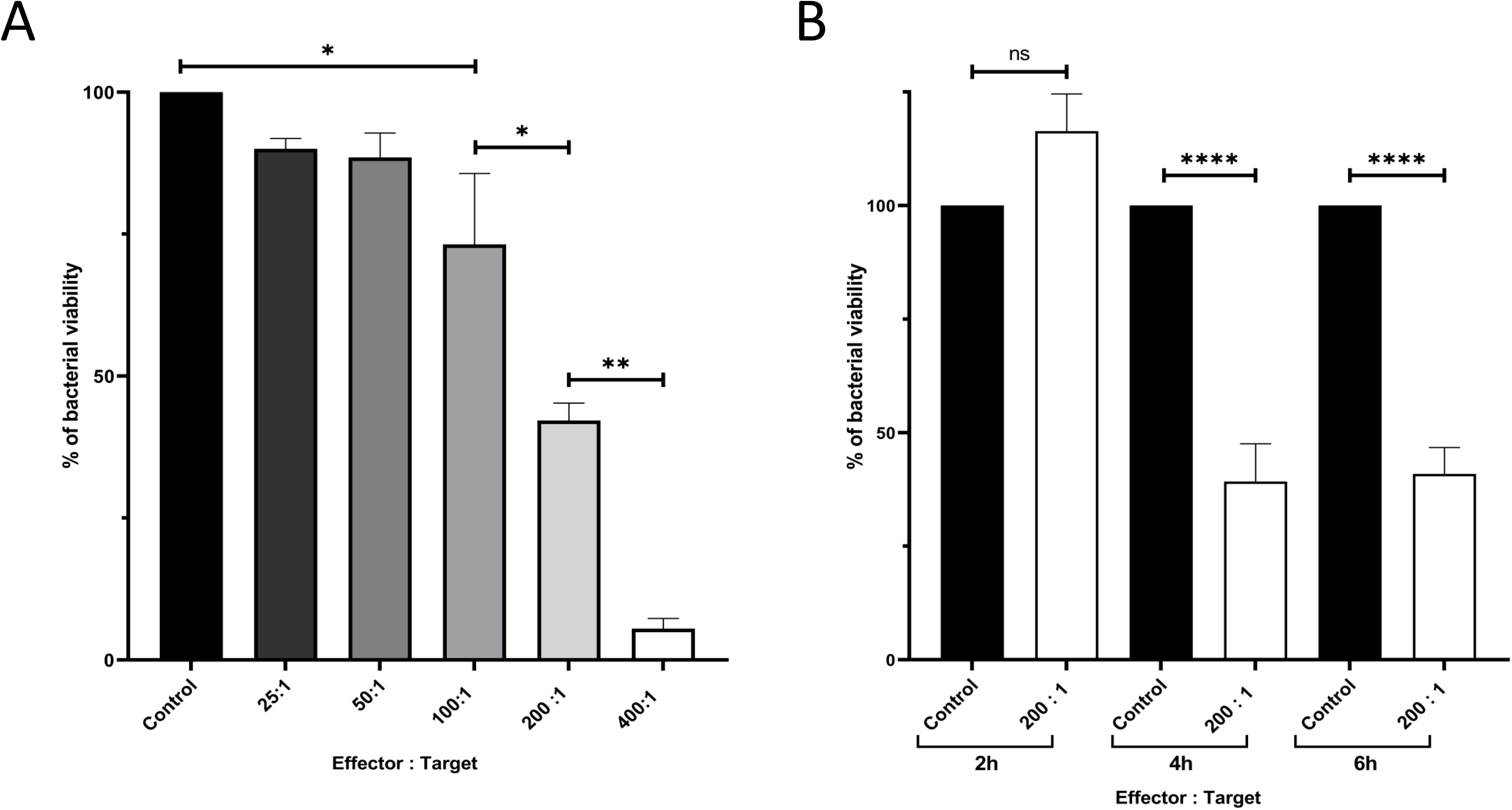
NK cells have an effect on PA14 growth. A) Effect of NK cells from healthy donors on PA14 growth (CFU/mL) after 6 h of co-incubation as a function of E:T ratio; B) For an E:T = 200:1, study over co-culture time of NK cells on PA14 (control = PA14 alone). CFU was determined by plate counting. These data were pooled from independent replicates using NK cells from 4 different donors carried-out (n= 4 and n=6, for the ratio and the kinetic studies, respectively). ** P≤0.01, **** = P≤0.0001. ns: not significant.

These results confirm that human NK cells impair *P. aeruginosa* growth *in vitro*, even if the bacterial strain is hypervirulent and known to induce NK apoptosis under certain conditions.

### NK cells from healthy donors have an activity against *S. aureus* bacteria

We then investigated whether this effect could be observed for a species quite different from *P. aeruginosa* also belonging to the ESKAPE group, i.e. *S. aureus*. Similar co-incubation experiments were performed to study the effect of NK cells on *S. aureus* growth for increasing E:T ratios and co-incubation time (figure 3). Remarkably, an E:T ratio of 0.5 is already sufficient to have a significant effect on *S. aureus* growth, reflecting a significantly higher efficacy than against PA14 (figure 2). The time for an optimal effect was observed after 6h of co-incubation.

**Figure 3:**
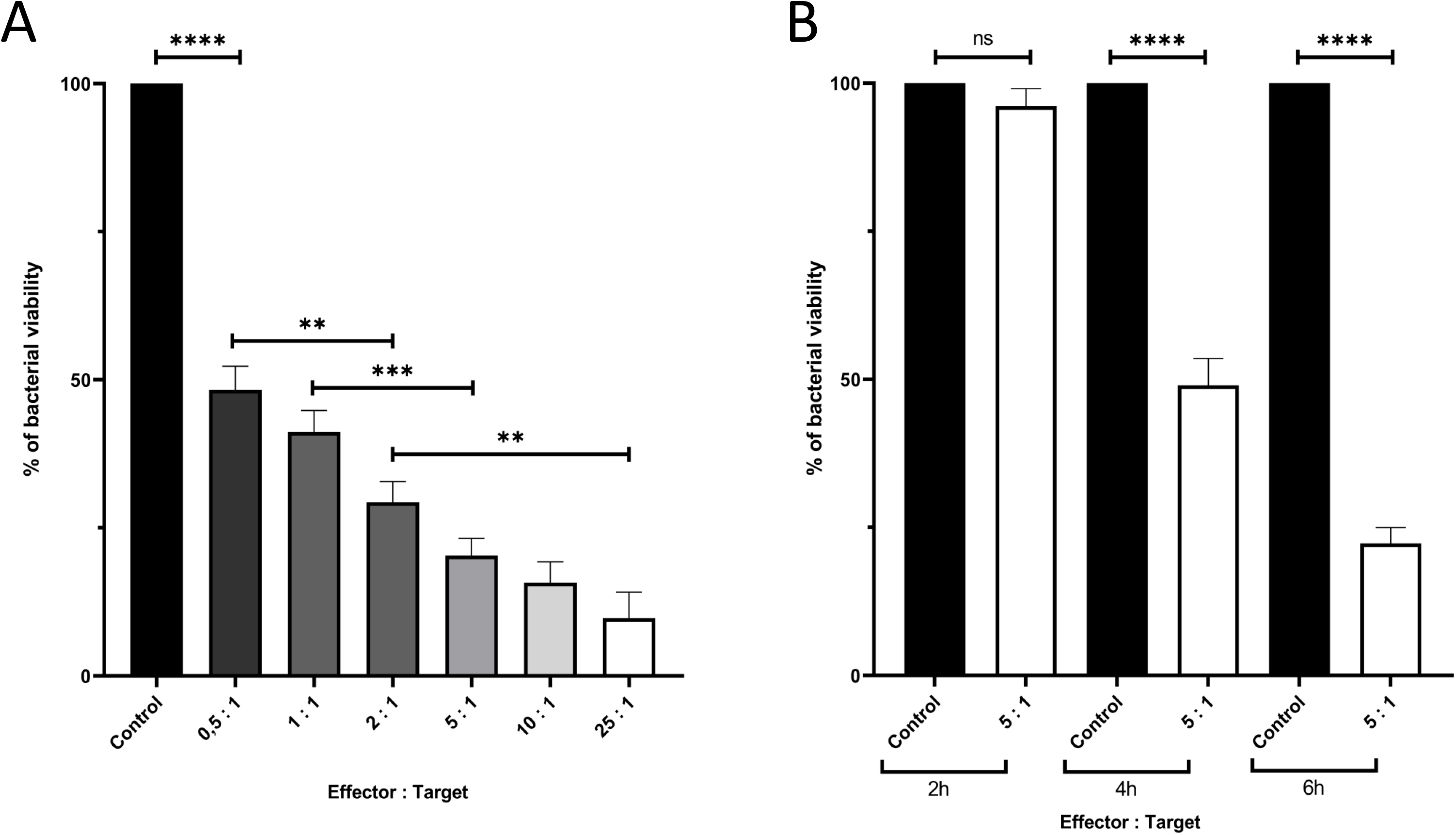
Effect of NK cells from healthy donors on *S. aureus* growth. A) Effect of different E:T ratios on the *S. aureus* Newman strain growth (CFU/mL) after 6 h of co-incubation; B) Study over co-culture time of the effect of NK cells on *S. aureus* CFU numbers (control = *S. aureus* alone); E:T = 5:1). CFU was determined by plate counting. These data were obtained from independent replicates (n= 7 for the E:T ratio study; n = 6 for the kinetic study) carried-out on different days using NK cells from 4 different donors. * P≤0.5, **** P≤0.0001. ns: not significant.

### NK92 cells also inhibit the growth of *S. aureus* bacteria

Having shown that NK cells from healthy donors impaired the growth of *S. aureus* growth, we sought to determine if the NK from a cell line (NK92-CD16) could also kill these bacteria. Indeed, the purity of NK cells from donors, although high (figure 1), is not 100% and the antibacterial activities we observed could in fact be the result of contaminating accessory cells. As shown in figure 4, NK92-CD16 cells killed *S. aureus* in a similar way (E:T ratio and time), demonstrating this effect was solely induced by NK cells, *i.e.* without the help of eventual contaminant accessory cells.

**Figure 4:**
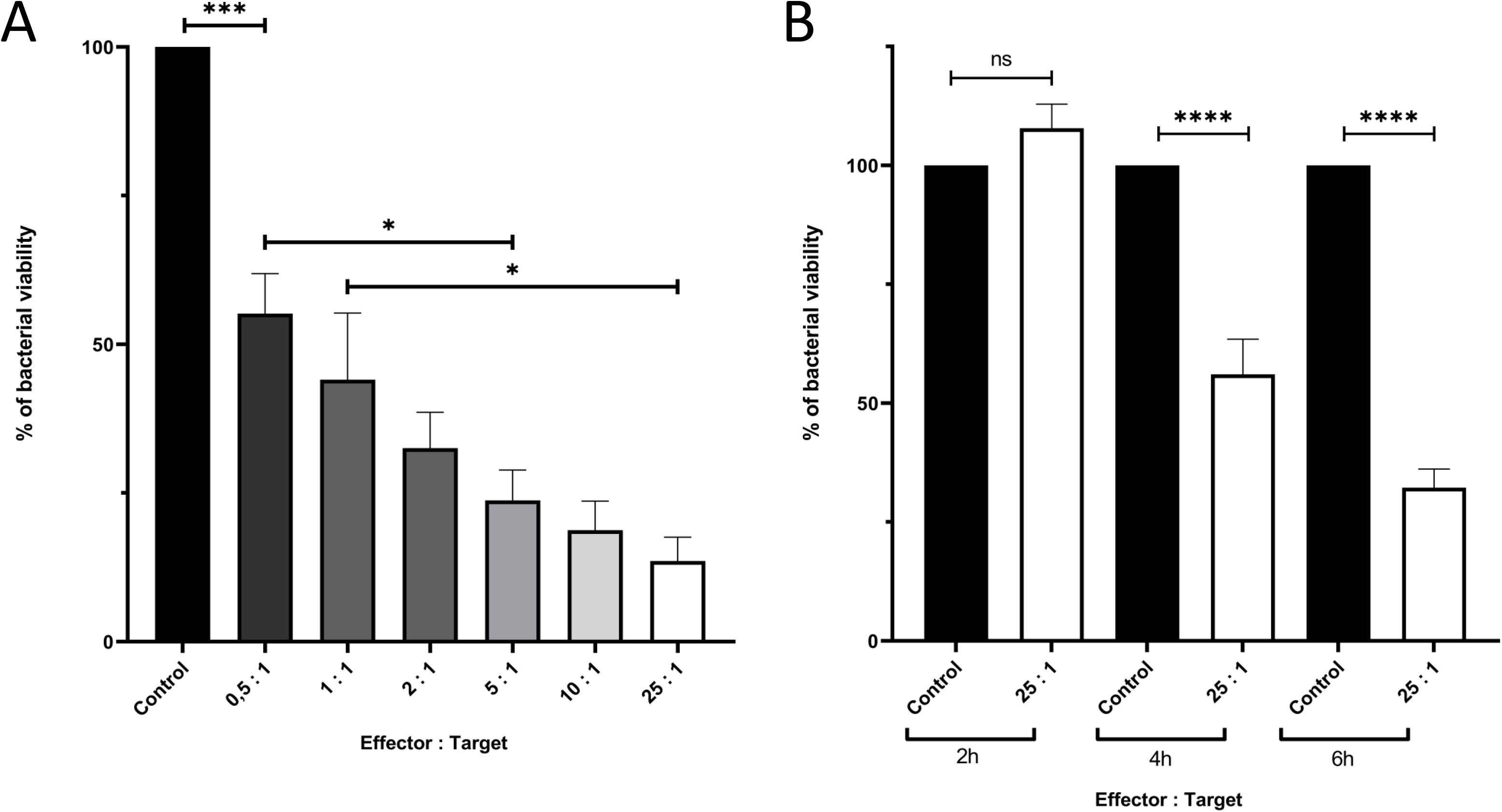
Effect of NK92-CD16 cells on *S. aureus* growth. A) Effect of different E:T ratios on the *S. aureus* Newman strain growth (CFU/mL) after 6 h of co-incubation; B) Study over co-culture time of the effect of NK cells on *S. aureus* CFU numbers (control = *S. aureus* alone); E:T = 25:1). CFU were determined by plate counting. These data were obtained from independent replicates (n= 5 for the E:T ratio study; n = 6 for the kinetic study) carried-out on different days using NK cells from 4 different donors. * P≤0.5, **** P≤0.0001. ns: not significant.

### Assessment of NK cell degranulation

These results led us to study whether this effect on bacteria growth is mediated by degranulation of NK cells. To address this point, we measured the ability of NK cells to degranulate upon contact with bacteria, through CD107a expression analyzed by flow cytometry. As shown on figure 5, for E:T ratio up to 100:1, *P. aeruginosa* bacteria did not induce expression of the degranulation marker CD107a. In contrast, *S. aureus* bacteria were able to induce the expression of CD107a even for an E:T ratio as low as 0.1:1 (figure 5). At an E:T = 1:1, up to ∼8.3 % of NK cells were stimulated by *S. aureus* bacteria for the expression of CD107a, reflecting an important level of degranulation.

**Figure 5:**
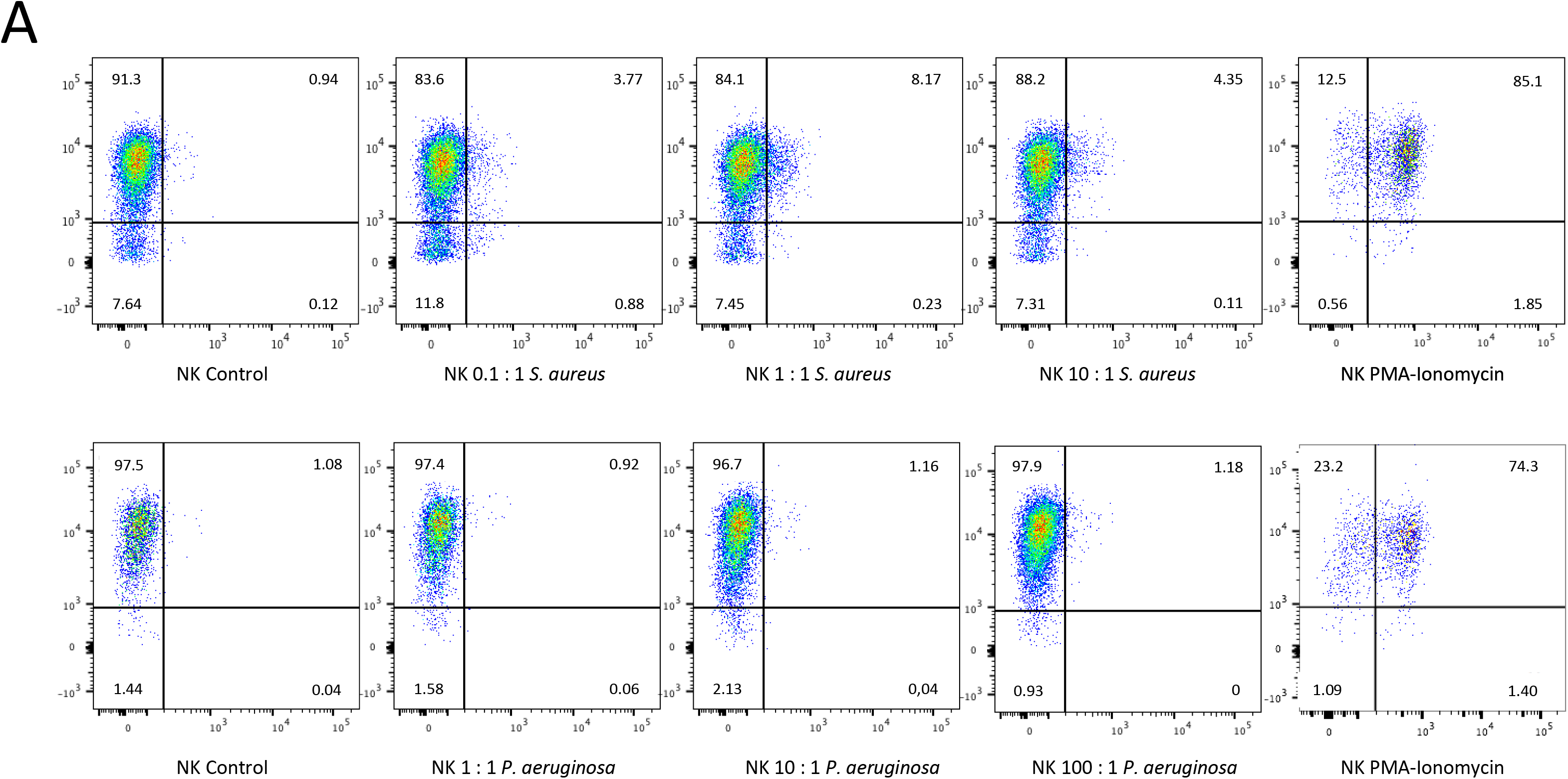
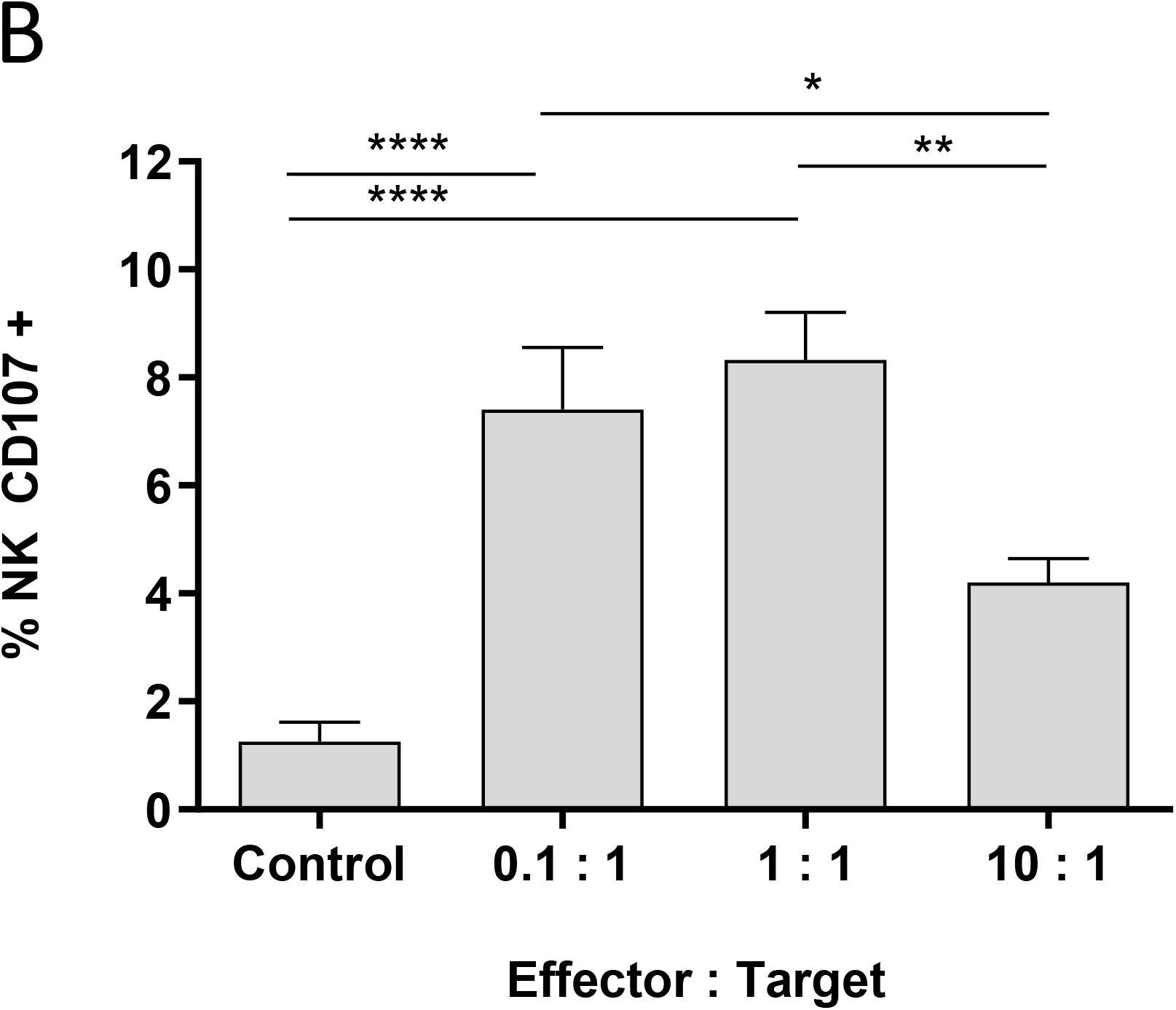
Measurement of CD107a expression on NK cells by flow cytometry. A) NK cells from healthy donors were co-incubated with PA14 *P. aeruginosa* or Newman *S. aureus* bacteria for 6 h at different E:T ratios and were stained with AF647-conjugated anti-CD107a and with PE-conjugated anti-CD56. The dot plots for CD107 and CD56-expressing NK cells are presented. B) The percentages of degranulation of NK cells from four donors for different NK:*S. aureus* ratios as a function of their potential of degranulation (CD107a+ PMA/Ionomycin) (n = 4). Degranulation controls were performed with NK cells stimulated with PMA/Ionomycin before analyzing samples by flow cytometry. * P≤0.1, ** P≤0.01, **** P≤0.0001.

## DISCUSSION

While NK cells are traditionally associated with viral and tumour responses, emerging evidences highlight their importance in combating bacterial infections. Thus, it was commonly accepted that NKs can inactivate bacteria, but mainly indirectly and with the help of accessory cells. However, several studies demonstrated that NK cells have a direct anti-microbial activity against extracellular *Salmonella typhi, Escherichia coli, Staphylococcus epidermidis, Burkholderia cepacia*, *Mycobacterium kansasii* and *Mycobacterium tuberculosis* bacteria, and [9–12]. In the most recent study, the group of Mody et al. reported this direct effect for two clinical isolates of *P. aeruginosa* and for the lab-strain PAO1 [13]. However, the clinical isolate, CF5, showed a different behaviour, since it was able to multiply better in the presence of NK cells than in their absence. Thus, we wondered how general these observations about a direct effect of NK cells on bacteria could be.

To this end, we first chose to test *P. aeruginosa* strain PA14. PAO1 and PA14 are two strains that were originally isolated from clinical samples and are now widely used in the laboratory to study *P. aeruginosa*. Interestingly, the PAO1 strain displays a moderate level of virulence, while PA14 is described as an hypervirulent strain [26]. This difference is due to several factors including the existence of two pathogenicity islands which encode for effectors such as ExoU via the type III secretion system (T3SS), a potent cytotoxin that induces cell damage and death. This strain also produces a higher level of pyocyanin than PAO1 [28], a toxin that generates reactive oxygen species (ROS) within host cells, leading to oxidative stress and damage to cellular components [29]. It has been reported that pyocyanin attracts neutrophils and induces their apoptosis [30] and that PA14 strain also induces death of NK cells, while a pyocyanin-deficient PA14 strain does not show any cytotoxic effect against NK cells [27]. However, *P. aeruginosa* K (PAK) has a cytotoxic effect on NK cells although it produces pyocyanin at low level, suggesting that other bacterial compounds are involved [27].

Interestingly, our co-incubation experiments show that NK cells from four donors are able to interfere with PA14 cultures (figure 2). However, large E:T ratios are necessary to decrease significantly (at least E:T = 200), or to prevent growth (E:T = 400) of bacteria after 6 h of co-incubation. This effect is notably more visible over time of co-incubation and is maximal in our setup at 4 hours. These results are very similar to those observed by the group of Mody et al for PAO1 strain and two other clinical strains (S35004 and CF127) [13]. It should be noted that after 24 h of co-culture, an effect of NKs against mycobacteria was already observed for an E:T = 20:1, but a higher ratio (E:T = 100:1) was also necessary to kill them in proportions similar to those for *P. aeruginosa* [9]. While to our knowledge the levels of pyocyanin productions for S35004 and CF127 strains are not described in the literature. Given our results for PA14 and those for PAO1 [13], it seems that the pyocyanin factor is not sufficient to explain why some strains of *P. aeruginosa* overcome the action of NK cells. The study of Chung et al. [27] aimed to understand the cytotoxicity mechanisms mediated by *P. aeruginosa*, thus they carried-out co-incubations experiments with E:T ratios of 0.1:1 and 1:1. The quite high E:T ratios used in our study and in that of the study of Mody’s group necessary to show an antibacterial effect are probably required to overcome the cytotoxic effect of *P. aeruginosa* strains against NK cells which is not due to factors specific to PA14 as observed by Chung *et al.* Interestingly, it has been shown that a contact between NK cells and yeast for at least 90 minutes is necessary to observe NK cell-induced cytotoxicity [31]. Unlike *S. aureus*, *P. aeruginosa* is mobile which could impede prolonged contact and explain why higher E:T ratios are required for *P. aeruginosa*. It will be interesting to further study the mechanisms used by *P. aeruginosa* strains to overcome the NK cell action in spite of quite high E:T ratios. *P. aeruginosa* being a gram-negative bacillus, we then wanted to investigate whether this effect of NK cells on extracellular bacteria could be observed for a quite different species that belongs to the ESKAPE group. Thus, we choose *S. aureus* as it is a Gram-positive coccus.

As shown in figure 3, a similar effect to that seen against *P. aeruginosa* could be observed for *S. aureus*, regardless of the donor from which the NK cells originate. However, an E:T ratio as low as 0.5 was already sufficient to get a 50% decrease in viable bacteria, while an E:T = 200:1 was necessary for PA14 strain to gain the same effect. This is also significantly lower than the minimal efficient E:T ratio of 12.5:1 reported for *Burkholderia cepacia* [10]. Our results demonstrate the powerful antibacterial activity of NK cells against *S. aureus*. Interestingly, lower but efficient ratio (E:T < 0.01) have been reported for purified NK from healthy donors against *Escherichia coli* [12]. Thus, it will be interesting to explore lower E:T ratios than those we have tested for *S. aureus*. As for *P. aeruginosa*, we found that this effect against *S. aureus* was maximal after 6 h of co-incubation. A previous work demonstrated that NK cells could be activated by *S. aureus* as shown by IFN-γ assays, but required a cell contact-dependent co-stimulation by monocytes [24]. FACS measurements on our NK cells preparations from donors excluded a contamination with monocytes. Furthermore, our results with NK92 cells exclude that NK cells need the help of any accessory cells to induce antibacterial activity (figure 4).

Previous studies have shown that the mechanism underlying the antibacterial activity of NK cells involves their degranulation, and the role of some of their granule components, such as perforin, granulysin and granzyme, have been demonstrated [9, 10, 13]. Surprisingly, our results show that NK cells degranulation behaviour differs depending on the bacteria used as targets. For *P. aeruginosa*, no degranulation was observed (figure 5A), while for *S. aureus* about up to 8.3 % of NK cells were degranulated in presence of bacteria (figure 5). The work of Feehan *et al.* demonstrated the role of both granzyme B and H contained in lytic granules for the killing of *P. aeruginosa*, by using inhibitor compounds and CRISPR Cas9-induced knockouts [13]. Considering that only quite high E:T ratios are efficient to induce an antibacterial effect, and that *P. aeruginosa* has been reported to induce a decrease of NKG2D expression thereby reducing NK cell cytotoxicity [32], it is possible that degranulation occurs at a such low level that it could be difficult to assess it with CD107a.

Overall, we confirm that NK cells without the help of accessory cells are able to mediate an antibacterial activity against *P. aeruginosa*, even though it is a hypervirulent strain, and we found that NK cells are also active against *S. aureus*, but in a much more efficient way. Future experiments should study the mechanisms by which NK cells are activated by *S. aureus* to exert their antibacterial activity and which NK effectors are crucial to this effect.

## CONCLUSIONS

This work reinforces previous studies showing NK cells can have an antibacterial role against various species of extracellular bacteria in the absence of accessory cells. How general in term of bacterial species sensitive to this effect will determine whether NK cells can be considered as a basis for the development of therapeutic options as an alternative or a complement to antibiotics. With antibiotic-resistant strains emerging worldwide, we believe that NK-based therapies could play a role, as they already do in combating tumour cells with the development of NK-cell engagers.

## ACKNOWLEDGEMENT

We thank the core facilitiy of SFR Santé: Cytometry and cell sorting “Cytocell”, for expert technical assistance. We are grateful to Alexis Broquet (Nantes, France) for the gift of the PA14 strain. B.L. received a fellowship from Région Pays-de-la-Loire and from AID (Agence de l’innovation de défense).

